# *Salmonella* Genomic Markers for Risk to Food Safety

**DOI:** 10.64898/2026.03.27.714810

**Authors:** Emma V. Waters, Claire Hill, Beata Orzechowska, Ryan Cook, Frieda Jorgensen, Marie Anne Chattaway, Gemma C. Langridge

## Abstract

Foodborne non-typhoidal *Salmonella* remains a major public health concern, yet many isolates recovered through food surveillance are not associated with human illness. To investigate whether genomic factors influence infection risk, we analysed whole-genome sequencing data from over 900 food and environmental isolates collected through UK Health Security Agency surveillance. Hierarchical clustering and comparative genomics identified distinct lineages associated with clinical cases, which were further contextualised using the global EnteroBase database. By combining pangenome and genome-wide association analyses, we identified distinct lineages within several serovars that differed in their association with human cases.

In *Salmonella* Agona, all clinical isolates belonged to a single lineage carrying a highly conserved 7 kb marker that was absent from low-risk strains and demonstrated strong sensitivity and specificity across global datasets. This marker was located within a prophage closely related to the well-characterised Fels-2 phage and encodes a DNA invertase previously implicated in phase variation, a mechanism that promotes bacterial adaptability. Our findings indicate that infection risk can be structured at the lineage level and associated with mobile genomic elements, particularly prophages, that may contribute to environmental persistence and host adaptation. This work advances genomic surveillance from retrospective linkage towards mechanistic and predictive risk assessment, with direct relevance for supporting risk-based decision-making during outbreak investigations.

**IMPORTANCE:** Not all strains found in food pose the same risk to human health, yet current surveillance systems generally treat them as equivalent hazards. We analysed over 900 genomes from food and environmental sources and found that human infection risk can be concentrated within specific genetic lineages rather than distributed across an entire serovar. In *Salmonella* Agona, we identified a highly conserved prophage-associated marker with strong sensitivity and specificity for infection-associated lineages. This marker was located within a prophage related to the well-characterised Fels-2 phage and encoded a DNA invertase previously associated to bacterial adaptation. These findings show how genome sequencing can move beyond outbreak detection to identify lineages with elevated public health relevance. Incorporating such information into surveillance programmes could improve risk-based decision-making, helping public health agencies prioritise investigations and interventions more effectively.

---

Infections caused by foodborne non-typhoidal *Salmonella* (NTS), typically resulting in self-limiting gastroenteritis, represent a significant public health and economic burden. Although most cases resolve without treatment, complications can require antimicrobial treatment, especially in vulnerable populations (1). Globally, NTS causes an estimated 93.8 million cases annually, with over 31,000 cases occurring in the UK and estimated to cost the economy £200 million annually (2, 3). Vigilant surveillance and timely detection and response to outbreaks are therefore effective crucial for infection control.

In England, the UK Health Security Agency (UKHSA) implemented routine Whole-Genome Sequencing (WGS) for *Salmonella* surveillance in 2015, substantially improving outbreak detection and source attribution through high-resolution genomic analysis (4–6). Nevertheless, delays can still occur between the onset of illness and the recognition of an outbreak, particularly when cases are sporadic, involve heterogenous bacterial populations or when food testing data is limited.

Beyond its established role in near real-time surveillance, WGS generates vast genomic datasets that are widely used for retrospective analyses, but are less routinely leveraged to inform predictive, risk-based prioritisation during future outbreak responses (7). These data offer the opportunity to better understand which *Salmonella* serovars, or even subtypes, are more likely to cause human disease, and which are not. While the presence of *Salmonella* in food is a public health concern, not all foodborne *Salmonella* isolates carry the same risk of causing illness and emerging evidence suggests that only a subset of serovars are responsible for the majority of clinical cases (8–10). If higher-risk foodborne *Salmonella* strains can be reliably identified, health agencies could prioritise interventions more efficiently and target their limited resources more effectively.

Our recent work explored this by examining the link between *Salmonella* isolates from food and those associated with human infection (7). By applying a data-driven approach to nearly 800 *Salmonella* isolates recovered from food and tested by UKHSA as part of routine surveillance and official control activities over a five-year period, we demonstrated that only a small proportion (∼10%) were genetically linked to clinical cases (7). This insight highlighted the potential to stratify foodborne *Salmonella* by risk and provides a basis for prioritising food testing and outbreak investigations (7). While this work demonstrated that the majority of foodborne *Salmonella* isolates do not contribute to human infection, it did not address whether specific genomic features could explain why certain strains pose a greater risk than others. Building on this foundation, here we applied a lineage-resolved genomic framework to investigate whether genetic variation within foodborne *Salmonella* serovars contributes to differences in their association with human infection. Across the 15 most common serovars, pangenome analysis and genome-wide association approaches identified lineages with differing associations with clinical cases. In *Salmonella enterica* serovar Agona (*S.* Agona), all clinical isolates belonged to a single cluster characterised by a highly conserved 7 kb genomic marker that demonstrated strong specificity and sensitivity for infection risk. The identification of this candidate marker supports a role for horizontally acquired elements in shaping infection risk and highlights their potential for predictive, risk-based surveillance.

## RESULTS

### Serovar distribution of food and environmental *Salmonella* in England

A total of 933 *Salmonella* isolates recovered from food (n = 799), environmental (n = 131) and pet food (n = 3) sources were referred to UKHSA’s Gastrointestinal Bacteria Reference Unit (GBRU) between January 2015 and December 2019 inclusive (Table S1) (7). Serotyping based on WGS data identified 87 different serovars in this collection (Table S2). However, the distribution was highly skewed with the 15 serovars most abundant accounting for 80.3% of all isolates and *S.* Heidelberg representing over one-third of the dataset (Tables 1 and S2).

**Table 1.** Distribution of foodborne and environmental *Salmonella* serovars across genetic clusters. Percentages of different *Salmonella* serovars in food, environmental and pet food samples referred to UKHSA’s GBRU from Food, Water and Environmental Microbiology Services (FWEMS) laboratories in England (2015–2019). This table shows the top 15 serovars accounts for 80.3% of all 933 isolates. The full distribution across all 87 different serovars in this collection is shown in Table S2. *Isolates classified as Typhimurium and Typhimurium Monophasic were combined due to one HC5 cluster containing both.

| Serovar | Number of isolates in original dataset | Percentage of original dataset (%) | Number of HC5 clusters in original dataset | Number of isolates in expanded dataset |
| --- | --- | --- | --- | --- |
| Heidelberg | 344 | 37.0 | 153 | 579 |
| Typhimurium | 56 | 6.0 | 18 | 2468* |
| Agona | 46 | 4.9 | 7 | 258 |
| Enteritidis | 46 | 4.9 | 17 | 8670 |
| Braenderup | 43 | 4.6 | 4 | 151 |
| Minnesota | 40 | 4.3 | 16 | 59 |
| Typhimurium Monophasic | 37 | 4.0 | 7 | 2468* |
| Infantis | 30 | 3.1 | 13 | 225 |
| Virchow | 20 | 2.2 | 10 | 66 |
| Brunei | 18 | 1.9 | 6 | 18 |
| Bareilly | 16 | 1.7 | 12 | 21 |
| Mbandaka | 15 | 1.6 | 8 | 22 |
| Weltevreden | 14 | 1.5 | 12 | 17 |
| Newport | 13 | 1.4 | 9 | 23 |
| Saintpaul | 11 | 1.2 | 7 | 26 |

### Hierarchical clustering reveals risk-associated lineages

Previous studies show a 5-SNP threshold is suitable for detecting clusters of closely related strains and is comparable in resolution to core genome Multi Locus Sequence Typing (cgMLST) (6, 11–14). Therefore, hierarchical clustering at the HC5 level (≤5 cgMLST allelic differences) was used to define genetically distinct groups (Table S3) (15). Per serovar, isolates falling into these HC5 clusters were searched for within the EnteroBase global *Salmonella* database to expand the initial dataset and provide a broader epidemiological context, increasing isolate numbers from 749 to 12,603 (Tables 1 and S3) (16).

To explore cluster associations with human infection, minimum spanning trees were generated for each of the 15 serovars, with isolates coloured by source metadata (Figs. 1 and S1) (17). Most serovars displayed limited separation between clinical and non-clinical isolates due to dominance of clinical isolates (*S*. Enteritidis and Typhimurium), lack of clinical isolates (*S*. Minnesota, Heidelberg and Brunei), or insufficient isolate numbers and were not analysed further.

**FIG 1.**
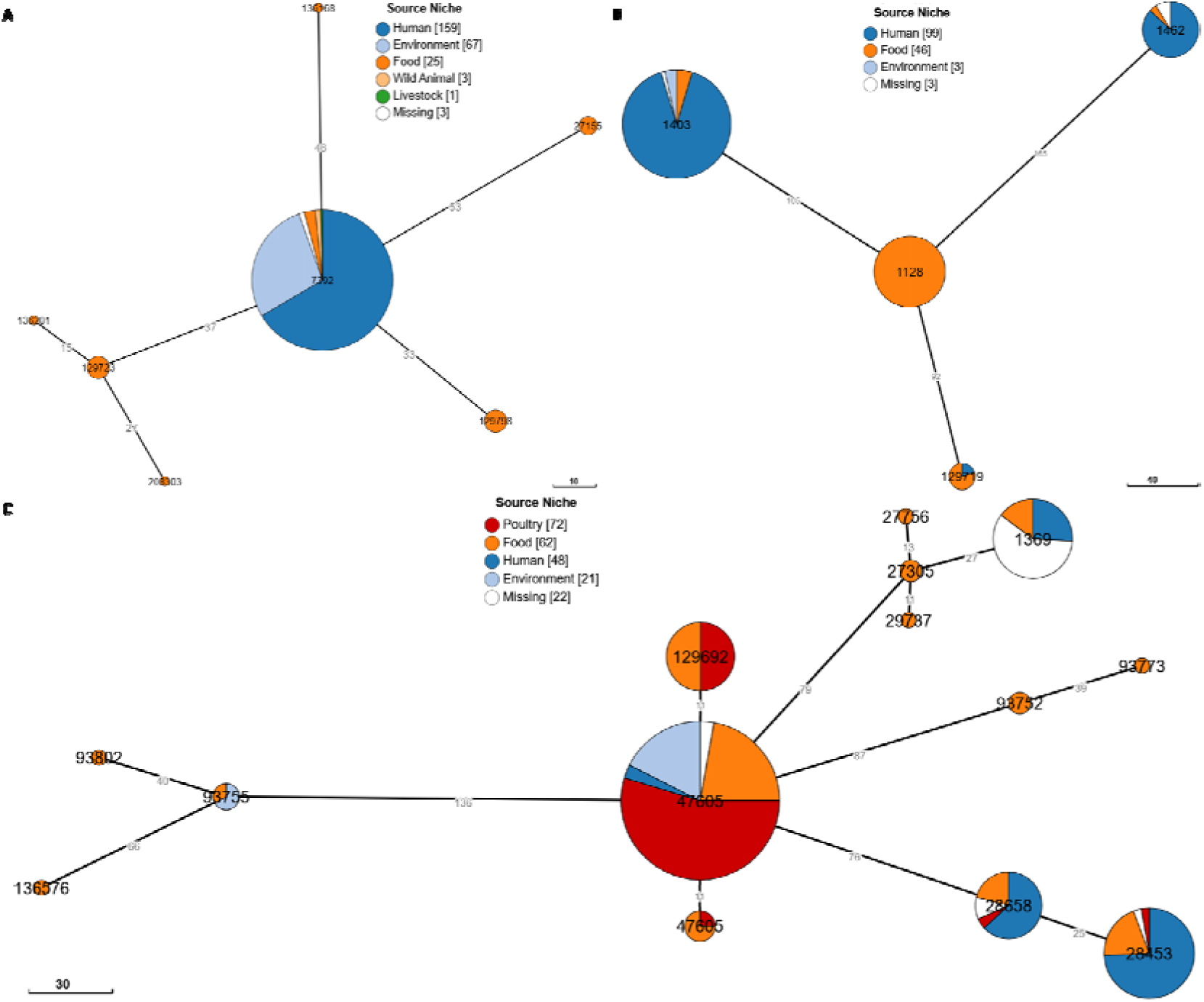
Source distribution within HC5 clusters of three *Salmonella* serovars showing distinct clinical and non-clinical clusters. Neighbour-joining cgMLST tree constructed using EnteroBase showing the presence of distinct HC5 clusters with and without associated clinical isolates in *Salmonella* serovars Agona (A), Braenderup (B) and Infantis (C). Circles are sized according to the number of isolates and coloured according to source: human (dark blue), environment (light blue), food (orange), wild animal (yellow), livestock (green), poultry (red) and missing (white). Data required to reproduce these figures is available via EnteroBase (Table S3). Scale bars represent the number of cgMLST allele differences which are shown on branches. HC5 IDs are given in bold inside circles.

In contrast, three serovars showed clear cluster segregation between clinical and non-clinical isolates, suggesting lineage-specific differences in propensity to cause human infection. The most striking pattern was observed for *S.* Agona, where all clinical isolates (159/258) were within a single HC5 cluster (HC5_7392) out of seven clusters in total (Fig. 1A). Based on this distinct separation, food and environmental isolates falling within the “risk” HC5 cluster that also contained clinical isolates were designated as “risk”-associated, while isolates belonging to “low-risk” clusters lacking clinical isolates were classified as “low-risk”. Using this definition, 239 isolates (159 clinical, 80 food/environmental) were designated as “risk” to human health, while 19 non-clinical isolates across the remaining six clusters were classified as “low-risk”.

Similar, though less pronounced, clustering patterns were observed for *S.* Braenderup (Fig. 1B) and *S.* Infantis (Fig. 1C). For *S.* Braenderup, none of the clinical isolates (99/151) were present in one HC5 cluster (HC5_1128), while the remaining three HC5 clusters contained clinical isolates with two of them (HC5_1403, 90% and HC5_1462, 87%) containing higher proportions of clinical isolates compared to the third (HC5_129719, 20%). For *S.* Infantis, all clinical isolates (48/225) were distributed across four of thirteen HC5 clusters (HC5_28453, HC5_28658, HC5_47605 and HC5_129692), suggesting that only specific lineages are associated with human infection (Fig. 1C).

To assess the strength of genomic links to human infection, we focused upon serovars *S.* Agona, Braenderup and Infantis based on their clinical relevance, representation across different sources, and the presence of distinct HC5 clusters with and without associated clinical isolates.

### Pangenome composition of specific serovars

Pangenomes were constructed for *S.* Agona, Braenderup and Infantis to explore genetic diversity and identify lineage-specific genes associated with risk (Fig. 2, Tables S4-5). *S.* Agona displayed the largest accessory genome, whereas *S*. Braenderup showed the most conserved pangenome but was enriched for prophage- and plasmid-associated genes.

**FIG 2.**
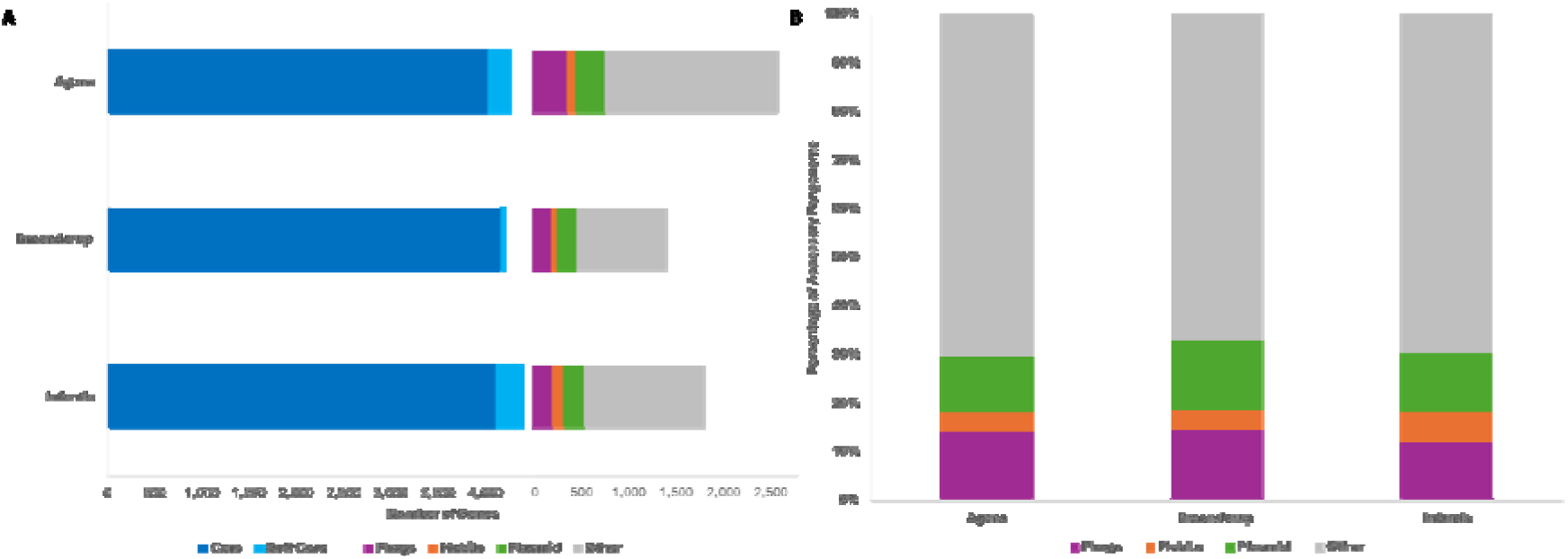
Distinct pangenome composition of three *Salmonella* serovars. (A) Stacked bar plot showing the composition of the pangenomes of *S.* Agona, Braenderup and Infantis, partitioned into core genes (blue, present in ≥99% of isolates), soft core genes (light blue, present in ≥95-99% of isolates), and accessory genes present in <95%. Bars represent raw gene counts. (B) Stacked column plot showing the proportional distribution of accessory gene categories within each serovar, expressed as a percentage of the total accessory genome. In both panels, accessory genes are further categorised by annotated function (see colour key).

### Genome-wide association identifies risk-associated genomic regions

To identify genomic features associated with increased and decreased likelihood of human infection (i.e. risk or low-risk markers, respectively), we used Scoary to assess associations between gene presence/absence and binary risk classification (risk vs. low-risk isolates) (18). In *S*. Agona and Braenderup, 344 and 154 genes, respectively, were significantly associated with either risk or low-risk clusters, while only one gene was identified in *S.* Infantis (Table S5). As expected, most significant genes (>99.3%) were in the accessory pangenomes.

Significant genes were mapped onto representative reference genomes to identify contiguous genomic regions of five genes or more (Table S5). In *S.* Braenderup, 89.6% (138/154) grouped into three contiguous genomic regions: a 63 kb region associated with increased risk (present in 77/111 isolates from risk clusters and absent from all low-risk isolates), and two smaller regions, measuring 44 and 8 kb, associated with reduced risk.

In *S.* Agona, 79.7% (274/344) of significant genes clustered into 13 contiguous genomic regions comprising five or more genes (Table S5). Two regions, measuring 26 and 7 kb, were associated with increased risk i.e. present in 237/239 isolates from the risk-associated HC5 cluster and absent from all low-risk isolates. The remaining 11 regions, ranging in size from 3-33 kb, were associated with reduced risk.

### Validation of candidate genomic markers in expanded datasets

Candidate genomic markers in *S*. Agona and Braenderup were validated using EnteroBase sequence data (16, 19). Marker sequences were used to generate locus-specific allele schemes and corresponding subtyping profiles (Table S6). Within expanded serovar datasets, these profiles showed strong concordance with HC5-defined risk classifications, with complexity generally increasing with marker length (Table S7). For example, the two *S*. Agona risk markers of 26 and 7 kb generated allele profiles containing 43 and 8 alleles, respectively. The 7 kb marker was highly conserved, with a single allele profile present in 237/239 risk-associated isolates and absent from all 19 low-risk isolates, corresponding to 99.2% sensitivity and 100% specificity (Table 8). The 26 kb marker displayed greater allelic diversity, generating 56 distinct allele profiles with partial allele presence in some isolates, but remained confined to risk-associated HC5 clusters (Table S7).

**Table 2.** Diagnostic performance of candidate *Salmonella* Agona genomic markers across validation datasets. Diagnostic characteristics of risk and low-risk genomic markers identified by genome-wide association analysis and validated using EnteroBase datasets of increasing epidemiological scale. Performance was evaluated in (i) the expanded dataset (n = 258) and (ii) a food-associated multi-serovar dataset comprising 12,827 isolates from the 15 most abundant foodborne serovars. For risk markers, the positive condition was defined as membership of HC5 clusters previously classified as risk-associated; for low-risk markers, the positive condition was defined as membership of low-risk HC5 clusters. Serovar spillover was assessed by quantifying marker presence in isolates from any of the other 15 top foodborne serovars. Full details are provided in Tables S7-8.

| Marker<br>(Risk Classification) | Dataset | Sensitivity<br>(%) | Specificity<br>(%) | Positive<br>Predictive<br>Value (%) | Negative<br>Predictive<br>Value (%) | Serovar<br>Spillover<br>observed? |
| --- | --- | --- | --- | --- | --- | --- |
| **SAL_XB6714AA_AS_**<br>**contig_18_42982_50209**<br>**- 7 genes 7 kb region**<br>**(Risk)** | Expanded | 99.2 | 100 | 100 | 90.5 | No |
|  | Food-associated<br>multi-serovar | 92.6 | 100 | 100 | 99.9 |  |
| **SAL_CB9225AA_AS_**<br>**contig_11_**<br>**456249_465620 - 11**<br>**genes 9 kb region**<br>**(Low-risk)** | Expanded | 36.8 | 100 | 100 | 95.2 | Yes<br>in 550<br>Typhimurium |
|  | Food-associated<br>multi-serovar | 2.7 | 95.6 | 1.3 | 98.0 |  |
| **SAL_JB0008AA_AS_**<br>**contig_35_22772_34010**<br>**- 17 genes 11 kb region**<br>**(Low-risk)** | Expanded | 36.8 | 100 | 100 | 95.2 | No |
|  | Food-associated<br>multi-serovar | 2.7 | 100 | 100 | 98.0 |  |
| **SAL_JB0008AA_AS_**<br>**contig_29_1_3788**<br>**- 4 genes 4 kb region**<br>**(Low-risk)** | Expanded | 36.8 | 100 | 100 | 95.2 | No |
|  | Food-associated<br>multi-serovar | 2.7 | 100 | 100 | 98.0 |  |
| **SAL_JB0008AA_AS_**<br>**contig_28_6118_13531**<br>**- 9 genes 7 kb region**<br>**(Low-risk)** | Expanded | 31.6 | 100 | 100 | 95.2 | Yes<br>in 4<br>Typhimurium |
|  | Food-associated<br>multi-serovar | 2.3 | 100 | 100 | 98.0 |  |

Similar allelic profile complexities (> three allele profiles) were observed for all *S.* Braenderup markers and for seven of the *S*. Agona low-risk markers and were therefore excluded from further analysis. The remaining four *S*. Agona low-risk markers (4-11 kb) generated one or three allele profiles unique to their low-risk HC5 clusters. Low-risk markers were present in six or seven of the 19 low-risk isolates and absent from all risk-associated isolates, corresponding to 31.6/36.8% sensitivity and 100% specificity (Table 8). Thus, while low-risk markers were highly specific, they captured only a subset of low-risk lineages.

### EnteroBase-wide validation of candidate genomic markers

Marker performance was further assessed using all 6,604 *S.* Agona isolates in EnteroBase (Table S7) (16, 19). The 7 kb marker remained strongly associated with risk-associated HC5 clusters (Fig. S2). Among risk-definable isolates (n = 6,079), it was present in 562/4,554 isolates from risk-associated HC5 clusters and rare in low-risk isolates (9/1,525), yielding high specificity (99.4%) and positive predictive value (98.4%) (Table S7). Across 12,827 isolates from the 15 most abundant foodborne serovars identified by UKHSA (Tables 1 and S8), the 7 kb risk marker was absent outside *S*. Agona, indicating strong serovar specificity (100%, sensitivity 92.6%).

In contrast, low-risk markers remained generally specific, but showed low sensitivity and variable positive predictive strength, reflecting their association with lineages infrequently linked to clinical disease and the clinical sampling bias inherent to EnteroBase. Limited cross-serovar presence was observed for two markers: the 9 and 7 kb markers were detected in 550 or 4 *S.* Typhimurium isolates, respectively. The other two low-risk markers remain restricted to *S*. Agona (Table S8).

### Genomic context and phage associations of marker regions

Having established the robustness, sensitivity and specificity of the five *S.* Agona genomic markers, we next investigated their genomic context and biological characteristics. Specifically, we assessed whether marker regions were conserved in genomic location, whether they were embedded within mobile genetic elements, and/or flanked by genes that might provide insight into functional roles in modulating infection risk.

Because EnteroBase assemblies are derived from short-read data, low-risk marker-containing contigs were identified using pangenome outputs and annotated assemblies to assess gene contiguity and conservation of gene order prior to analysis of flanking regions. From our previous work we had access to hybrid assemblies of 14/237 risk marker containing isolates and used these to investigate the genomic context of the risk marker (20).

All markers, irrespective of risk classification, contained prophage-associated genes and were located within larger prophage regions, with the exception of the 4 kb low-risk marker (Table S9). For all but one of the four low-risk markers, short-read contigs did not extend beyond the marker or the surrounding prophage region on both sides, preventing detailed assessment of flanking genomic context (Table S9). Where flanking sequences were available across multiple isolates, low-risk markers were generally located within conserved chromosomal regions.

Comparison of flanking sequences revealed that four of the five markers, irrespective of risk classification, occurred within the same ∼102 kb chromosomal region encompassing 50 genes. Within this locus, 27/28 isolates investigated contained either: 1) the 7 kb risk marker; 2) the 9 kb low-risk marker; or 3) both the 4 kb and 11 kb low-risk markers together. This suggests that this region of the *S.* Agona chromosome represents a prophage integration hotspot where alternative prophage configurations occur.

This 102 kb prophage hotspot was enriched for genes involved with core cellular functions, membrane biogenesis, stress response, transport and host interaction. Flanking genes systems involved in acid tolerance (*cadABC*), iron acquisition (*fepA, fes, iroBC*) and antimicrobial peptide resistance (*mig-14, virK*), and the type III secretion system effector *pipB2*.

In one isolate (SAL_TB1655AA_AS), the 11 kb low-risk prophage was located outside of this hotspot, surrounded by genes associated with surface structures (including fimbriae, pili and adhesins), environmental sensing, and mobile genetic elements. These flanking regions included several insertion sequence elements and transposases, as well as toxin-antitoxin system components (*symE* and Ibs family toxin), suggesting local genomic plasticity. For the 7 kb low-risk marker-containing contigs, only one of six isolates contained sufficient flanking sequence that extended beyond the prophage region to determine genomic location, which was located outside the hotspot.

### Characterisation of candidate prophage risk marker

The 7 kb risk marker was prioritised for further investigation due to its a strong association with clinical risk-associated lineages. High-quality hybrid assemblies from representative isolates enabled detailed genomic characterisation of the region (20).

Analysis indicated this marker formed part of a ∼34 kb prophage with strong sequence similarity (∼97%) to *S.* Typhimurium LT2 phage Fels-2 (NC_010463) (21), with 35/43 (76.1%) predicted proteins showing homology (Fig. 3, Table S9). Further inspection revealed that the previously identified 26 kb risk marker and the 7 kb marker formed different halves of the same ∼ 34 kb prophage, with the 7 kb marker localised to one end of the prophage and predominantly encoded tail-associated proteins together with a DNA invertase (Fig. 3, Table S9). DNA invertases are site-specific recombinases implicated in phase variation, surface antigen expression, and adaptive phenotypes in enteric bacteria, including modulation of host interaction and immune evasion (22, 23). Together, these features suggest that the marker may influence phenotypes relevant to host interaction and infection risk. Targeted deletion of either the 7 kb region or the invertase gene produced no detectable effect in motility and *Galleria mellonella* infection assays (Figs. S3-4 and Supplementary Material).

**FIG 3.**
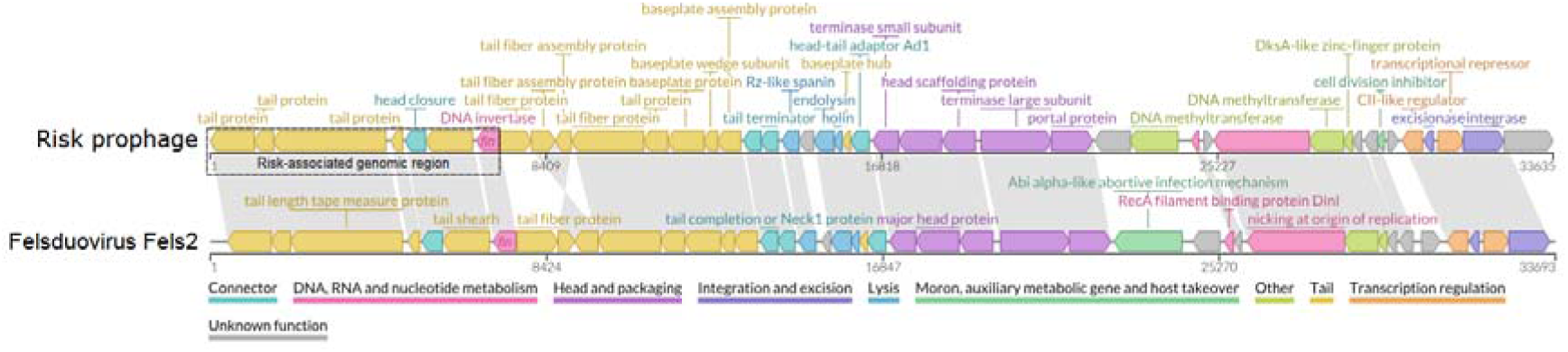
Genomic organisation of the prophage containing the 7 kb risk marker compared to its closest relative, *Salmonella* phage Fels-2. Conserved genes are indicated by grey shading, and genes are coloured according to their predicted functional categories (colour key). The 7 kb risk marker region is outlined and contains the *fin*-like DNA invertase gene.

## DISCUSSION

Our previous work showed that most foodborne *Salmonella* isolates do not contribute to human infection, highlighting the challenge of identifying which strains pose the greatest public health risk (7). Although over 2,500 *Salmonella* serovars have been described globally (24), the population circulating in England is far less diverse. In this study, only 87 serovars were identified, with epidemiological burden further concentrated within a small subset, as 15 serovars accounted for >80% of isolates. This highly skewed distribution emphasises the importance of risk-based surveillance and prioritisation of control efforts toward those of highest risk.

However, serovar abundance in surveillance datasets does not necessarily reflect true prevalence in food or disease risk. For example, *S.* Heidelberg was highly represented, largely reflecting targeted surveillance with UKHSA’s nationwide monitoring of *Salmonella* in imported food, including poultry products samples at the point of import from Brazil (25), rather than true prevalence in food. While all foodborne *Salmonella* isolates undergo routine WGS, many originate from reactive sampling prompted by outbreaks, complaints, or follow-up investigations after previous detection. Consequently, food surveillance datasets inevitably contain elements of sampling bias. These discrepancies underscore the limitations of serovar-level classification for assessing public health risk and highlights the need for higher-resolution genomic approaches capable of distinguishing lineages with differing infection risk.

Hierarchical clustering revealed that lineages within serovars varied considerably in their relationship to human infection. In *S.* Enteritidis and Typhimurium, clinical and non-clinical isolates were intermixed across HC5 clusters, suggesting broadly distributed risk across genetically diverse lineages. This may reflect extensive genetic diversity of these serovars, driven by horizontal gene transfer or repeated introductions from geographically distinct sources (26, 27).

In contrast, serovars such as *S.* Minnesota, Brunei and Heidelberg were frequently detected in food or environmental samples but rarely associated with human infection, indicating limited capacity to cause disease. For example, despite the reported presence of *S*. Heidelberg and Minnesota in imported Brazilian poultry (28–30), their contribution to human illness in England appears minimal (25). Similarly, national surveillance data indicate that between 2015 and 2019 only 0.38%, 0.08% and 0.02% of 37,673 clinical salmonellosis cases were attributed to *S.* Heidelberg, Minnesota and Brunei, respectively (7). This discrepancy likely reflects a combination of factors, including effective detection and removal of contaminated consignments at import, the further processing of the contaminated poultry which sufficiently removes *Salmonella* before retail, or intrinsic differences in virulence and infectious dose among serovars (7).

A different pattern was observed for *S.* Agona, Braenderup and Infantis, where infection was associated with specific HC5 clusters. The most striking example was *S.* Agona, where all clinical isolates belonged to a single HC5 cluster, suggesting a lineage with enhanced association with human infection. Similar, although less pronounced, patterns were observed for *S.* Braenderup and Infantis. Together, these findings indicate that for some serovars infection risk can be concentrated within specific genomic lineages rather than uniformly distributed across a serovar. This observation provides a strong rationale for investigating the genetic determinants that distinguish these lineages and may contribute to differences in their propensity to cause human disease.

Comparison of pangenomes across the three focal serovars revealed differences in accessory genome structure. *S.* Agona exhibited the largest accessory genome, consistent with greater genomic heterogeneity, although it may also be influenced by the larger number of isolates included in this pangenome analysis. In contrast, *S.* Braenderup had a more conserved genome, suggesting lower overall genomic plasticity, but contained the highest proportion of prophage- (14.4%) and plasmid-associated (14.5%) genes, highlighting the importance of mobile genetic elements in shaping diversity within this serovar. A similar proportion of plasmid-associated genes was observed in *S.* Agona (14.2%), although prophage-associated genes were less common (11.7%). Genes linked to other mobile elements occurred at comparable levels in both of these serovars (3.6–4.0%) but were more prevalent in *S.* Infantis (6.4%).

It is important to note that as these pangenomes were constructed from isolates in food-associated HC5 clusters, they reflect diversity within circulating food-associated populations rather than the global serovar-level diversity. Nevertheless, the high proportion of accessory genes associated with mobile elements across all three serovars supports a central role for horizontal gene transfer in lineage differentiation within foodborne *Salmonella*. In particular, prophage-associated genes appear to contribute substantially to accessory variation and may contribute to phenotypic differences between lineages. This interpretation is consistent with our subsequent genome-wide association analyses, which identified several lineage-associated genomic regions embedded within prophage elements, including the risk-associated markers identified in *S.* Agona. Prophages are well-recognised drivers of bacterial evolution and virulence in *Salmonella*, often introducing genes involved in host interaction, stress tolerance and regulatory variation (31–33).

Genome-wide association analysis further supports this, with the vast majority of significant genes (>99%) belonging to the accessory pangenomes and largely clustered into discrete contiguous genomic regions (>79%) rather than occurring independently. In *S.* Agona, two regions (26 and 7 kb) were strongly associated with risk-associated lineages, while 11 genomic regions were associated to low-risk clusters but typically occurred only in subsets of low-risk isolates. Interestingly, ten of these regions were shared among the same small group of six low-risk isolates, suggesting either co-acquisition through horizontal gene transfer or subsequent retention within a distinct lineage that has evolved separately from the high-risk cluster.

Only one significant gene association was identified for *S.* Infantis, potentially reflecting lower statistical power, a more complex genetic architecture underlying infection risk, or polygenic determinants that are not captured by presence-absence genome-wide association approaches.

Within the expanded dataset, marker-specific allele schemes demonstrated strong concordance with HC5-defined risk classifications. The 7 kb *S.* Agona marker showed near-perfect performance (99.2% sensitivity, 100% specificity), supporting its use as a genomic indicator of risk-associated lineages. A second 26 kb marker was also confined to risk lineages but exhibited greater allelic complexity, likely reflecting subsequent variation following initial acquisition. In contrast, low-risk markers were highly specific (100%) but showed substantially lower sensitivity (31.6-36.8%), capturing only subsets of low-risk clusters. This asymmetry suggests that increased infection risk may be driven by acquisition of a conserved genomic element, whereas low-risk phenotypes may reflect greater genetic heterogeneity.

Evaluation across EnteroBase confirmed that the 7 kb risk marker retained strong predictive performance at global scale and remains entirely restricted to *S.* Agona. This strong within-serovar specificity and consistent association with clinically relevant lineages supports its value as a stable genomic indicator of infection risk. On the other hand, low-risk markers showed markedly lower performance at global scale (0.5–1.0% sensitivity, 6.1–90% PPV), reflecting their presence in only a small subset of low-risk lineages and the inherent clinical sampling bias present within EnteroBase. Cross-serovar spillover was also observed for some low-risk markers, including detection within *S.* Typhimurium isolates, suggesting that certain prophage-associated elements may retain some capacity for horizontal mobility across serovars.

For four of five markers, despite differing risk classifications, localised to a shared ∼102 kb chromosomal region, consistent with a prophage integration hotspot in *S.* Agona. Isolates typically contained either the 7 kb risk-associated marker or alternative low-risk markers, indicating mutually exclusive prophage configurations. In this context, differences in infection risk may therefore reflect the identity of the prophage occupying this site, rather than the simple presence or absence of accessory genes. This hotspot lies within a conserved genomic neighbourhood enriched for genes involved in environmental stress response, membrane function and host interaction. Integration at this site may influence neighbouring gene expression or host-adaptation pathways, suggest that the broader genomic context may also contribute to lineage-associated differences in infection risk.

Interestingly, the highly conserved 7 kb marker formed part of the larger 34 kb Fels-2-like prophage, the remainder of which corresponded to the more variable 26 kb risk marker. While the 26 kb region accumulated substantial allelic diversity across risk-associated isolates (56 allele profiles), the 7 kb region remained completely conserved (1 allele profile), suggesting stronger evolutionary constraint and explaining its superior performance as a marker of infection-associated lineages.

Despite the strong epidemiological association, deletion of the 7 kb marker region or its DNA invertase did not affect motility, or virulence in *Galleria mellonella*, indicating it is not a standalone virulence determinant. Instead, it may contribute to infection-associated lineage success through context-dependent regulatory or ecological effects not captured in these assays.

An additional layer of interpretation arises from the identified DNA invertase showing homology to the Fels-2-associated recombinase Fin. In *Salmonella*, phase variation of flagellin expression, specifically switching between the alternative flagellins FliC and FljB, is a well-characterised mechanism that generates phenotypic heterogeneity (34), enabling immune evasion and adaptation to fluctuating host environments (35). This process is classically controlled by the Hin invertase; however, Fin has been shown to catalyse inversion of the same H-segment and can function redundantly when Hin is impaired (23). Experimental studies, primarily exclusively in *S.* Typhimurium, have demonstrated that while single *hin* or *fin* mutants retain the ability to switch flagellar phase, loss of both recombinases abolishes phase variation entirely (23). However, as *S.* Agona are monophasic and lack the classical flagellar phase variation system, this invertase likely acts on alternative targets. Its association with infection-linked lineages suggests that invertase-mediated regulatory systems encoded by prophages may contribute to lineage success and host adaption more than previously appreciated. In this regard, our genome-wide association approach has effectively rediscovered a known adaptive mechanism, within a different serovar and in association with epidemiologically defined risk. Notably, the distribution of Fels-2-like elements across isolate collections suggests that their presence may not be required for strains already adapted to clinical infection but instead may confer an advantage during transition between environmental or food-associated niches and the human host. Consistent with this, only a minority of isolates (32/208, 15.4%) in our broader *S.* Agona clinical collection carried *fin*-like DNA invertase (20), and similar trends have been reported elsewhere, where Fels-2-like prophages were not significantly associated with disease (42/54, 77.8% of isolates from infected humans, animals or food) but were present in all environmental isolates (n = 17) (36). Together, this supports a model in which Fels-2 contributes to ecological flexibility and transmission potential, rather than acting as a direct determinant of virulence.

Overall, these findings support a model in which infection risk is shaped by lineage-specific genomic configurations, particularly alternative prophage elements occupying conserved chromosomal hotspots. These elements may act as stable markers of high-risk lineages, even where direct functional effects are subtle, and illustrate the value of genomic analysis with epidemiological data to advance the development of predictive, lineage-informed approaches to foodborne risk assessment.

## MATERIALS AND METHODS

### Hierarchical clustering and dataset expansion

Raw WGS data were uploaded and assembled on EnteroBase as previously described (20). Sample data was collated using accession numbers - all assembled samples available as of 13/1/25 were included for dataset expansion.

HierCC (v1) and cgMLST (v2) were used to assign isolates to HC5 clusters (groups of closely related strains, differing by ≤5 cgMLST alleles) (15). For the 15 most common serovars, all HC5 clusters were identified and used to retrieve additional isolates from EnteroBase to provide international epidemiological context. Minimum spanning trees were generated using MSTree (v2) and visualised on GrapeTree, with isolates coloured by source metadata to facilitate comparison of clusters with risk status (16, 17).

### Pangenome analysis and genome-wide association analysis

The open platform Galaxy (v23.0.6) was used to perform subsequent bioinformatic analyses. Genome assemblies for *S.* Agona, Braenderup and Infantis were downloaded from EnteroBase, annotated using Bakta (v1.9.3) with the genus flag *Salmonella* (37), and used to construct serovar-specific pangenomes with Roary (v.3.13.0) using default parameters (38). These pangenomes are available on request. Accessory genes were functionally categorised using keyword-based searches (e.g. “phage”, “plasmid”, “transposase”, etc).

HC5 clusters were trait-classified as “risk” (≥1 clinical isolate) or “low-risk” (non-clinical only), and this trait-classification was then applied to all isolates within each cluster. Gene-risk associations were evaluated using Scoary (v1.6.16) (18), with statistical significance determined following Bonferroni correction with adjusted *p*-values < 0.05 (Table S5).

### Investigation of genomic context of statistically significant genes

Bakta-generated gene IDs were collected for statistically significant genes from representative reference genomes to identify contiguous genomic regions (≥5 genes). For *S*. Braenderup, two reference genomes were used: the food isolate SAL_KA8987AA_AS (HC5_1128) and the human isolate SAL_BB3835AA_AS (HC5_1403). For *S*. Agona, four reference genomes were used: three food isolates (SAL_BA8032AA_AS, HC5_27155; SAL_CB9225AA_AS, HC5_129798; SAL_JB0008AA_AS, HC5_129723) and the human isolate SAL_XB6714AA_AS (HC5_7392).

### Validation of sensitivity and specificity of genomic markers

Marker performance was evaluated using EnteroBase data - all samples of assembled status as of 1/10/25 were included to generate these further datasets as described below.

Marker sequences were used to perform a locus search within EnteroBase, generating locus-specific allele schemes and corresponding allele profiles in which numerical allele identifiers represented marker presence, while absence was recorded as zero values (Table S6) (16, 19). These, in combination with HC5 cluster information, were applied to three datasets to assess marker performance at increasing epidemiological scale: 1) expanded datasets (*S*. Braenderup, n = 148; *S.* Agona, n = 258); 2) all *S.* Agona isolates in EnteroBase (n = 6,604); and 3) a food-associated, multi-serovar dataset (n = 12,827 isolates from the 15 most abundant serovars) (Tables S7-8).

Markers generating >3 allele profiles within their expanded dataset were excluded from further analysis. Sensitivity, specificity, and positive predictive values were calculated from confusion matrices based on HC5-defined risk classification. Within the global *S.* Agona dataset, diagnostic characteristics were calculated using only HC5 clusters with risk-definable isolates (n = 6,079). Cross-serovar presence was assessed in the multi-serovar dataset.

### Genomic context and phage analysis of marker regions

For low-risk marker-containing isolates where only short-read assemblies were available, marker-containing contigs were identified from annotated assemblies and used to extract up to 100 upstream and downstream genes (Table S9). For the 7 kb risk marker, annotated hybrid assemblies (n = 14) were used construct a pangenome before extracting 100 upstream and downstream genes (20, 38) (Table S9). Gene content was aligned across all marker-positive isolates to assess conservation and infer the genomic location (Table S9).

For all markers, prophage content was confirmed using Proksee, Phigaro and PHASTEST (39–41). For risk isolates 112 and 126, prophages were predicted using geNomad (v1.11.1) and compared to the INPHARED database (November 2025) using MASH (v2.3) (42–44). The risk-associated prophage was annotated using Pharokka (v1.8.2) using -g prodigal for gene calling and compared to the *S.* Typhimurium LT2 prophage Felsduovirus Fels-2 (accession: NC_010463) genome using LoVis4u (v0.1.6) to assess synteny (45, 46).

## ACKNOWLEDGEMENTS

EVW and GCL gratefully acknowledge the support of the BBSRC Institute Strategic Programme Microbes in Food Safety BB/X011011/1 and their constituent project /E/QU/230002B. CH is funded through the BBSRC grant Exploring variation within *Salmonella* Dublin using sequencing and multi-model phenotyping to define markers of virulence and zoonosis BB/Z516673/1. RC is funded through the BBSRC grant Bacteriophages in Gut Health BB/W015706/1.

This study is funded by the NIHR Health Protection Research Unit (NIHR200892) in Genomics and Enabling Data at University of Warwick in partnership with UKHSA, in collaboration with University of Cambridge and Oxford. MAC is based at UKHSA. The views expressed are those of the author(s) and not necessarily those of the NIHR, the Department of Health and Social Care or UKHSA.

The authors would like to thank Leanne Sims, Marker Webber and Jemma Snell for their assistance with setting up the *Galleria* model.

## AUTHOR CONTRIBUTIONS

GCL and MAC designed the methodology and conceived the study. EVW and BO curated the data. EVW and CH performed validation and investigation. EVW, CH and RC performed formal analysis and visualisation. EVW and GCL wrote the original draft of the manuscript. All authors reviewed and edited the manuscript.

