## Supplementary Material for "*Salmonella* Genomic Markers for Risk to Food Safety"

**SUPPLEMENTARY MATERIALS**

**Table S1** List of 933 isolates recovered from food (n = 799), environmental (n = 131) and pet food (n = 3) sources referred to UKHSA’s GBRU from FWEMS laboratories in England between January 2015 and December 2019 inclusive.

**Table S2** Prevalence of different *Salmonella* serovars in food, environmental and pet food samples referred to UKHSA’s GBRU from FWEMS laboratories in England (2015 - 2019).

**Table S3** List of isolates included within the expanded dataset with each tab representing one of the 15 most prevalent *Salmonella* serovars. Typhimurium monophasic isolates were included alongside those identified as Typhimurium.

**Table S4** Functional association of accessory genes from pangenomes of all isolates of *Salmonella* Agona, Braenderup and Infantis associated with food- and environmental-related isolates.

**Table S5** Scoary results for associations between gene presence/absence from risk classification (risk versus low-risk HC5 clusters) across the pangenomes of all isolates of *Salmonella* Agona, Braenderup and Infantis associated with food- and environmental-related isolates.

**Table S6** EnteroBase allele schemes defining candidate genomic marker subtyping. Each tab corresponds to a single genomic marker identified through genome-wide association analysis and summarises the locus identifiers, positions and function description provided by Bakta and EnteroBase.

**Table S7** EnteroBase-wide validation of candidate *Salmonella* Agona and Braenderup genomic markers subtyping schemes associated with risk or low-risk of human infection. Each tab corresponds to a single genomic marker identified through genome-wide association analysis. Within each tab, marker performance is evaluated across up to three datasets of increasing epidemiological scale to assess sensitivity and specificity as detailed in the methods.

**Table S8** Presence or absence of the five candidate *Salmonella* Agona genomic markers across a food-associated dataset of 12,827 isolates representing the 15 most common foodborne serovars identified in England.

**Table S9** Genomic context of the *Salmonella* Agona risk and low-risk markers region in representative hybrid assemblies and short-read contigs, respectively. Marker regions are highlighted in light green. Conserved genes shared between assemblies are highlighted in dark green. Genes located within prophage regions predicted by PHASTEST are highlighted in blue.

| Strain | Knockout status | Source | Reference |
| --- | --- | --- | --- |
| *S*. Agona 112 | parent | UKHSA | (1) |
| *S*. Agona 126 | parent | UKHSA | (1) |
| *S*. Agona 112ΔInv | DNA invertase | This study |  |
| *S*. Agona 126ΔInv | DNA invertase | This study |  |
| *S*. Agona 112Δ7kb | 7 kb region | This study |  |
| *S*. Agona 126Δ7kb | 7 kb region | This study |  |

**Table S10** Strains used in this study.

| **Primer/Plasmid** | **Sequence/Source** | **Description** |
| --- | --- | --- |
| HR1_Inv_F | GGCTACGGTCTCACTACTA  GCGCTTTCATGCCCGTTTTC | Primers for the amplification of Homologous Region 1  of the DNA invertase identified in *S*. Agona 112 and 126 |
| HR1_Inv_R | GGCTACGGTCTCTCTCCAA  GTGGCGATATTGTACCAGC |  |
| HR1_7kb_F | GGCTACGGTCTCGCTACG  ATGTAGGAATTTCGGAC | Primers for the amplification of Homologous Region 1  of the 7 kb region identified in *S*. Agona 112 and 126 |
| HR1_7kb_R | GGCTACGGTCTCACTCCTT  GGTTCTCAATATTGAGTG |  |
| HR2_F | GGCTACGGTCTCTCGCTAA  CAGGCCCCCTCGCAAA | Primers for the amplification of Homologous Region 2  of both the DNA invertase and the 7 kb region  identified in *S*. Agona 112 and 126 |
| HR2_R | GGCTACGGTCTCATCGTTA  TTATCATTATTCTGACGGTG  AACCATTGG |  |
| pDOC-AmpR | (2, 3) | Gene doctoring base plasmid used for golden gate  construction of knockout plasmids |
| pDOC_Inv | This study | Generated for this study, golden gate assembly of HR1,  HR2, *kan*R and pDOC-AmpR to generate knockouts of  the DNA invertase region identified |
| pDOC_7kb | This study | Generated for this study, golden gate assembly of HR1,  HR2, *Kan*R and pDOC-AmpR to generate knockouts of  the 7 kb region identified |
| pEACH | (2, 3) | Donor plasmid for Kanamycin resistance casette (*kan*R)  flanked by FRT removal and BasI sites for golden gate  cloning |
| pACBSCE | (2, 3) | Helper plasmid containing the λ-Red genes for  recombination and I-SceI Endonuclease for plasmid  cleavage of pDOC and self |
| pCP20 | (4) | Temperature senitive *kan*R removal plasmid encoding  FLP recombinase |

**Table S11** Primers and plasmids used to make knockout strains in this study. The authors would like to thank Emma Holden and Mark Webber for providing the plasmids pDOC-AmpR, pEACH, pACBSCE and pCP20 used to generate genomic knockouts.

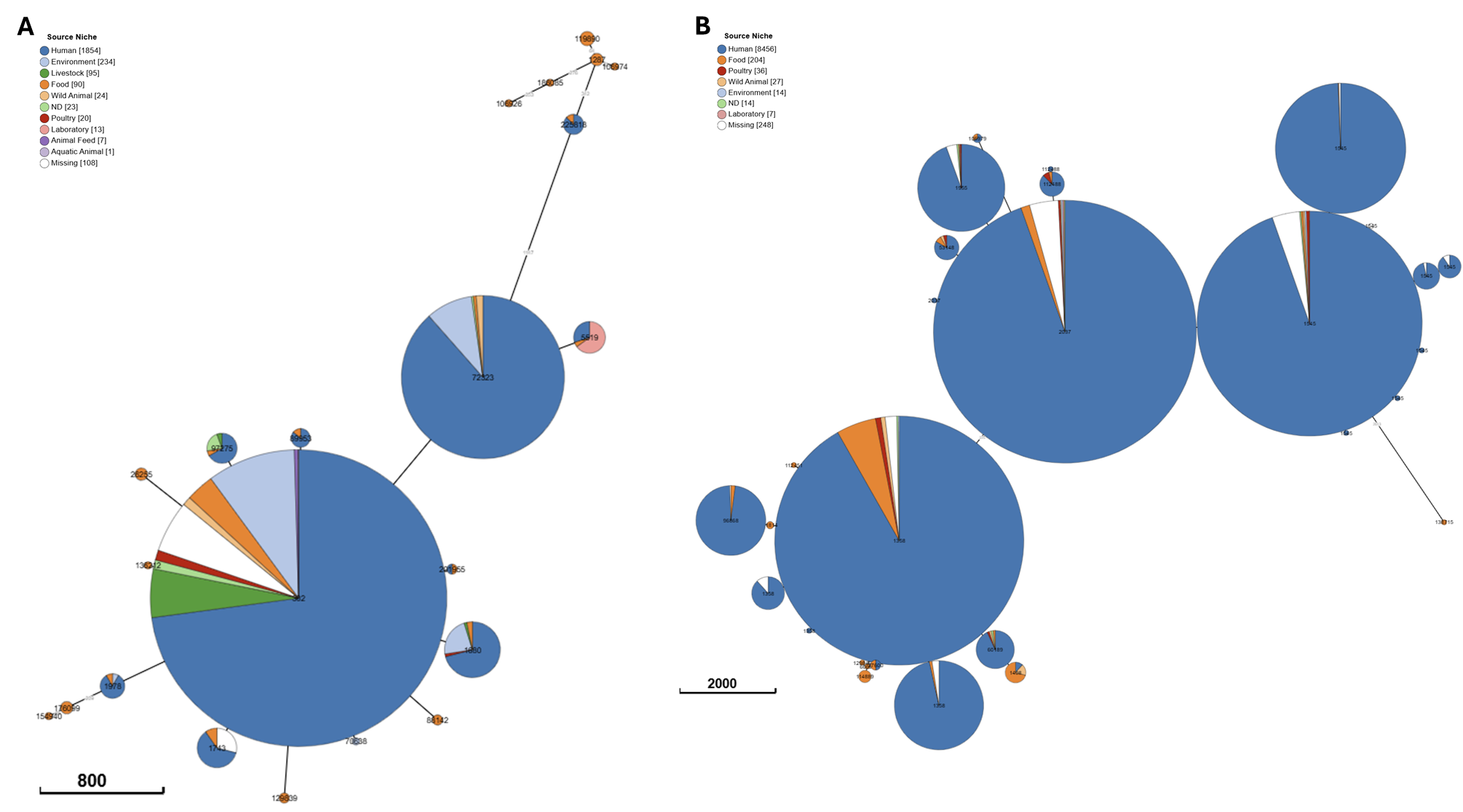

**FIG S1** Source distribution within HC5 clusters of two *Salmonella* serovars showing clinical isolates dominated all clusters. Neighbour-joining cgMLST tree constructed using EnteroBase showing the dominance of clinical isolates in *Salmonella* serovars Typhimurium* (A) and Enteritidis (B). Circles are coloured according to source niche as shown in keys. The data required to reproduce these figures is available via Enterobase (Table S3). Scale bars represent the number of cgMLST allele differences which are also shown on branches. *Typhimurium monophasic isolates were included alongside those identified as Typhimurium.

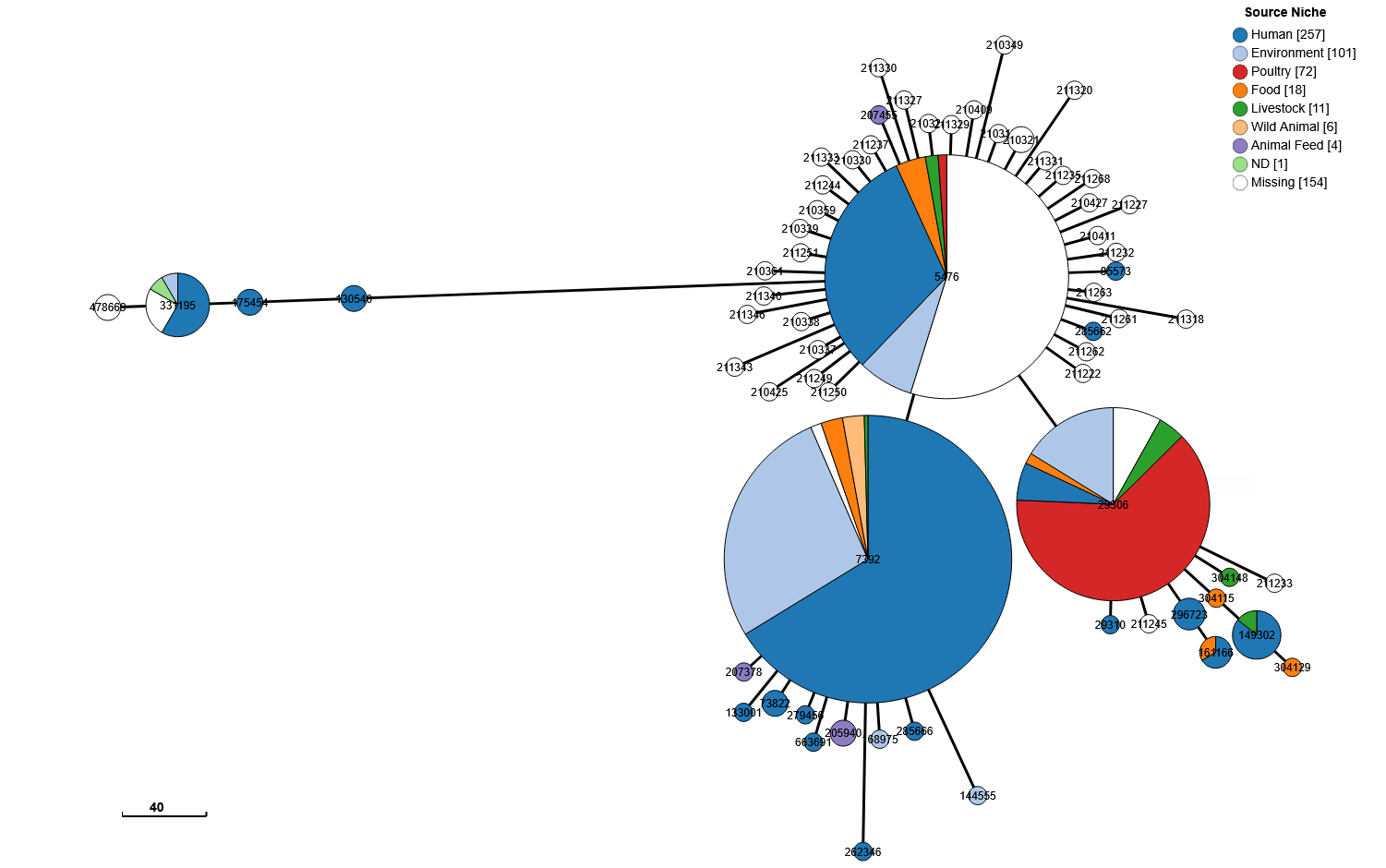

**FIG S2** Source distribution within 85 HC5 clusters containing the *Salmonella* Agona risk marker. Neighbour-joining cgMLST tree constructed using EnteroBase showing the food isolates cluster with clinical isolates. Circles are coloured according to source niche as shown in keys. The data required to reproduce these figures is available via Enterobase (Table S7). Scale bars represent the number of cgMLST allele differences which are also shown on branches.

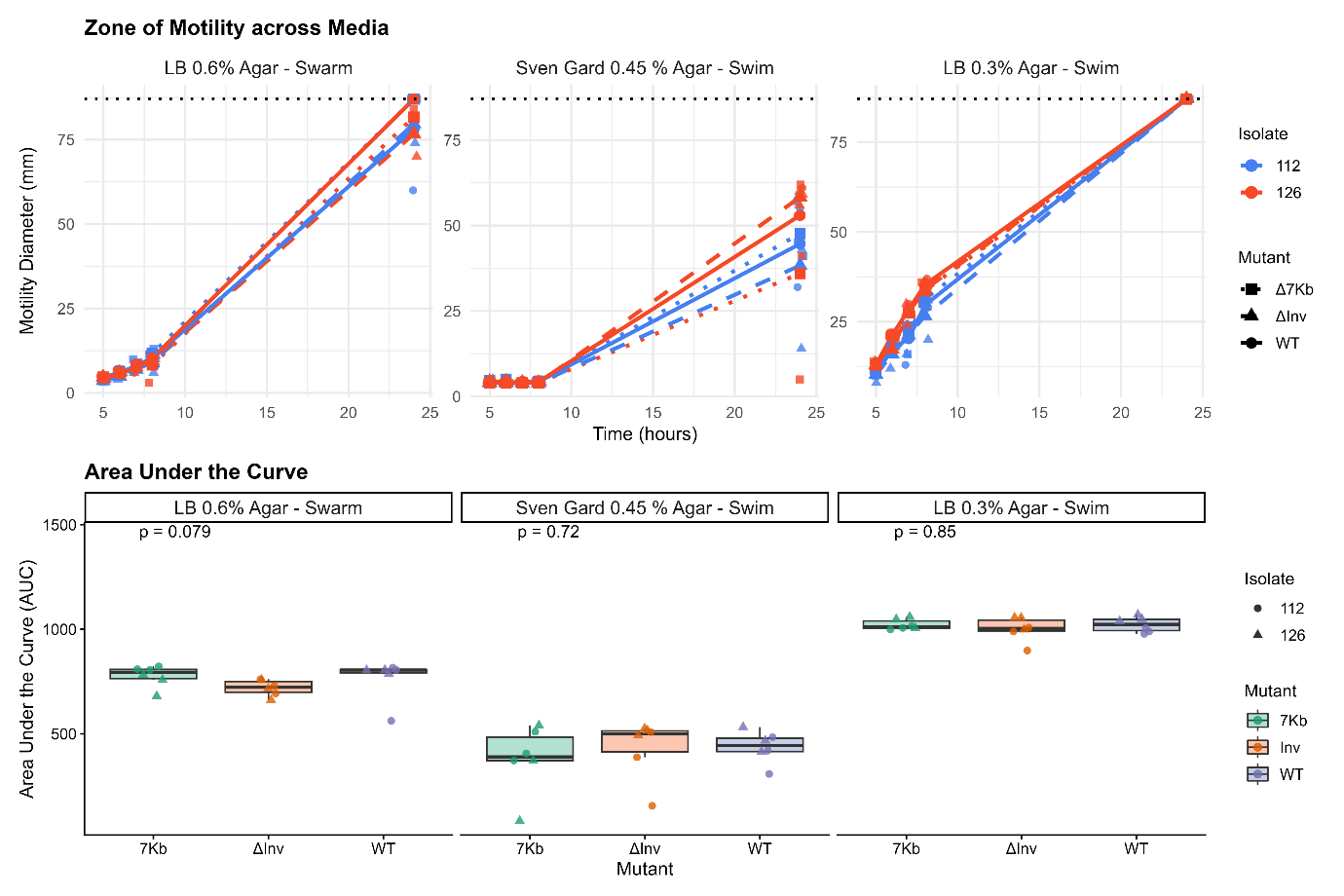

**FIG S3** Zones of motility of wildtype and mutants measured across 24 hours.

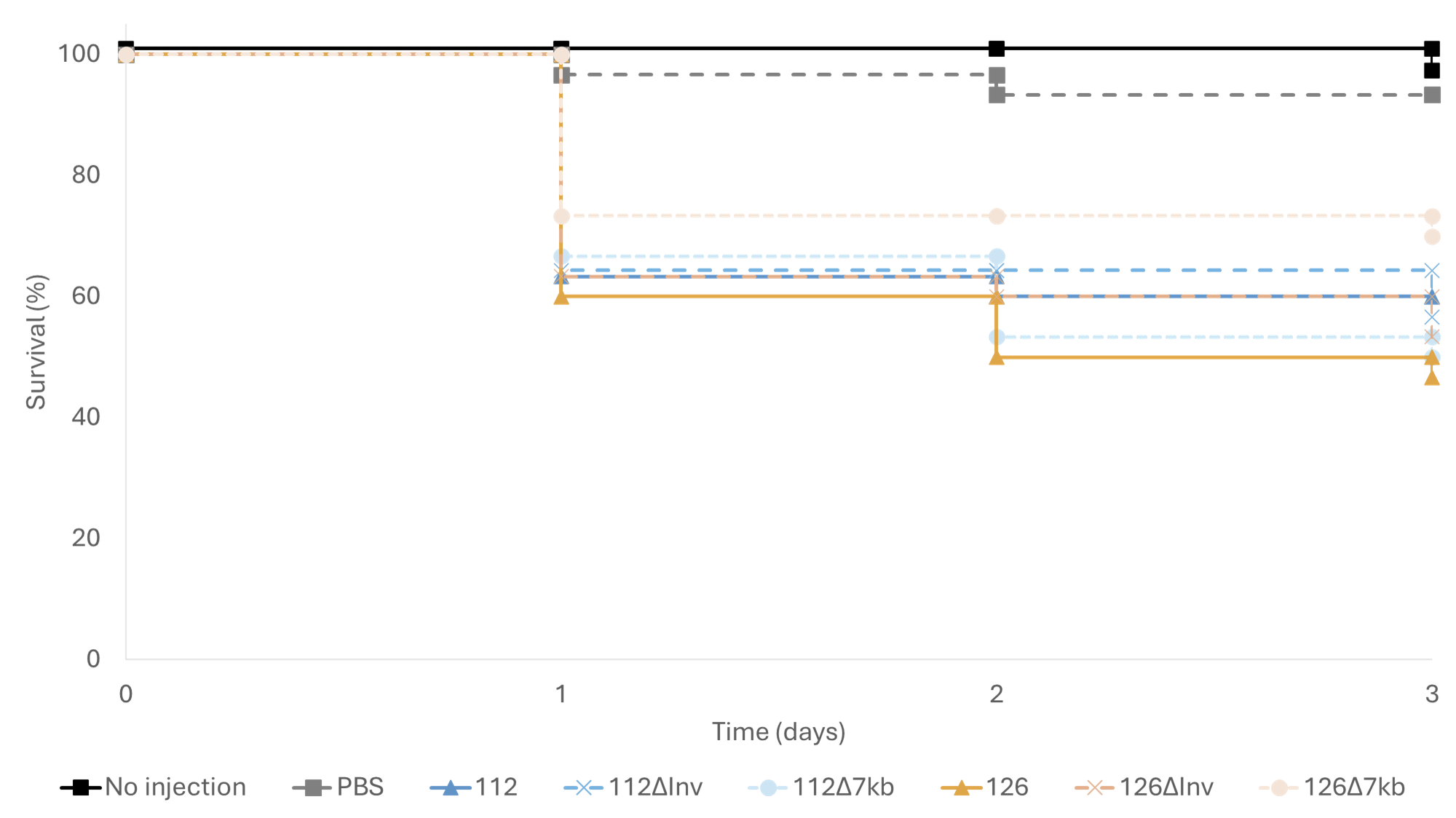

**FIG S4** Survival of *Galleria mellonella* larvae following infection with wild-type and knockout strains. Kaplan–Meier survival curves of larvae (n = 10 per condition, performed in triplicate) infected with 10⁵ CFU per larva. Knockout strains carried deletions of either the 7 kb marker region (Δ7kb) or the DNA invertase gene (ΔInv), with corresponding wild-type parental strains shown for comparison. Larvae injected with PBS alone served as negative controls. Survival was monitored over 3 days post-infection. Kruskal–Wallis tests of the area under the curves indicated no significant differences in survival between wild-type, ΔInv, and Δ7kb strains within each background (112: *p* = 1.00; 126: *p* = 0.43).

**SUPPLEMENTARY METHODS**

**Generation of genomic knockouts** All strains, plasmids and polymerase chain reaction (PCR) primers used in this study are listed in Tables S10-11. Knockouts of the 7 kb region and the DNA invertase gene were generated using gene doctoring (2, 3, 5). For each knockout, the 450 bp homologous regions immediately upstream (HR1) and downstream (HR2) of the target regions were amplified by high-fidelity PCR using Q5 Polymerase (New England Biolabs, USA) with a 5’ primer overhang containing BsaI restriction site. PCR products were size confirmed on 1% agarose gel and purified using a Zymoclean Gel DNA Recovery Kit (Zymo Research, USA) according to the manufacturer’s instructions.

Knockout plasmids pDOC_Inv and pDOC_7kb were constructed by Golden Gate assembly of pDOC-AmpR plasmid, the kanamycin resistance cassette flanked by BasI sites from the plasmid pEACH (3), HR2 DNA and the respective HR1 DNA. The assembled plasmids were transformed into competent *E*. *coli* DH5α (Invitrogen, USA) by heat shock at 42 °C for 45 s (6). Plasmids were purified using Zypppy Plasmid Miniprep kit (Zymo Research, USA) and verified by sequencing (Plasmidsaurus low-copy plasmid service, <25 kb).

*Salmonella* cells were made electrocompetent by growth in 2X Yeast Extract Tryptone medium to early exponential phase (OD_600_ 0.25-0.35), followed by preparation as described (7). Electroporations were performed as described previously (8). For chromosomal integration and gene deletion, *S.* Agona strains 112 and 126 were first transformed with the helper plasmid pACBSCE (2), encoding the λ-Red recombination system, and subsequently with the appropriate pDOC_7kb or pDOC_Inv knockout plasmid.

Chromosomal integrations were induced as described (2). Briefly, single colonies carrying both plasmids were grown in 500 μL Luria-Bertani (LB) broth at 37 °C for 3 hours. Cells were pelleted, washed three times, and resuspended in 500 μL 0.1x LB supplemented with 0.3% arabinose before incubation at 37 °C for 3 hours to induce recombination. 100 μL were plated on LB-no-salt agar supplemented with 50 mg/L kanamycin and 5% sucrose and incubated overnight at 37 °C.

The kanamycin resistance cassette, flanked by FLP recombination target (FRT) sites, was removed by transforming with pCP20, a temperature-sensitive plasmid encoding FLP recombinase and AmpR resistance gene (4). Carbenicillin-resistant transformants were selected at 30 °C. Successful transformants were then screened for loss of antibiotic resistances at 42 °C. Sensitivity to both kanamycin and carbenicillin indicated FRT removal and pCP20 loss, this was confirmed by PCR.

**Whole-genome sequencing and bioinformatic analysis to confirm knockouts** Genomic DNA from knockout mutants was extracted using Fire Monkey kits (RevoluGen, UK) according to the manufacturer’s instructions. DNA was normalised to 5ng/µL, and 2 µL was used for library preparation. Each sample was combined with 0.5 µL Tagmentation Buffer (Illumina, USA), 0.5 µL Bead-Linked Transposomes and 4 µL PCR-grade water, then incubated at 55 ⁰C for 10 mins. Following tagmentation, 10 µL KAPA 2G Fast Hot Start Ready Mix (Merck, Germany), 2 µL PCR-grade water and 1 µL of 10 µM primer mix containing both P7 and P5 Illumina barcodes was added (9). PCR amplification was performed with 72 ⁰C for 3 mins, 95 ⁰C for 1 min, 14 cycles of 95 ⁰C for 10 s, 55 ⁰C for 20 s and 72 ⁰C for 3 mins.

Libraries were quantified using the QuantiFluor® dsDNA System (Promega, USA) and run on a D5000 ScreenTape (Agilent, USA) using an Agilent Tapestation 4200 to confirm successful library preparation. Libraries were pooled in equal amounts, followed by double-SPRI size selection (0.5 and 0.7X bead ratios) using sample purification beads (Illumina, USA). The final pool was quantified and assessed using QuantiFluor® and D5000 ScreenTape to determine molarity. Sequencing was performed on an Illumina Nextseq2000 instrument following the manufacturer’s denaturation and loading protocols.

Raw reads were mapped to the parental hybrid genome sequence using BWA-MEM (v0.7.17.1) (10), before being visualised in the Integrative Genomics Viewer to confirm deletion of the target regions (11).

**Confirmation of fitness, virulence and motility of knockouts** Growth kinetics of parent and knockout strains were compared and confirmed to be indistinguishable by measuring optical density at OD_600_ at 10 mins intervals across 72 hours using a BioTek LogPhase 600 microplate reader (Agilent, USA) set at 37 °C, 500 rpm.

Relative virulence of parent and knockout strains was assessed using the *Galleria mellonella* infection model (12, 13). Larvae were chilled at 8 °C for 30 mins prior to injecting and inoculated in the second hindmost proleg with a 25 μL Hamilton syringe. For each isolate, ten larvae were injected with 10 μL of bacterial suspension containing 10^8^ CFU/mL. Two control groups were included: ten larvae injected with 10 μL PBS and ten non-injected larvae.

Following injection, larvae were incubated in the dark at 37 °C and mortality was recorded at 24, 48, and 72 hours post-infection. For each strain, data from at least three independent experiments were combined. Survival curves were generated for visualisation and differences in survival between groups were assessed using Kruskal–Wallis tests of the area under the curve.

The motility of parent and knockout strains was assessed using both swarming and swimming assays (14, 15). Overnight cultures were adjusted to 0.6 OD_600_ and 3 µL was spotted onto a 25 mL LB swarm agar plate (10g/L tryptone, 5g/L yeast extract, 10g/L NaCl, 0.6% agar and 0.5% final glucose) and a 25 mL Sven Gard agar plate (12.7g/L peptone, 1.2g/L yeast extract, 5g/L NaCl, 0.45% agar and 0.15% final glucose). The swimming assay was performed by piercing a 25mL LB swim agar plate (10g/L tryptone, 5g/L yeast extract, 10g/L NaCl, 0.3% agar) to half of the depth and inoculating with 3 µL of OD_600_ adjusted overnight cultures. Plates were air dried at room temperature for 10 mins and then incubated at 30 °C for 24 hours with the zones of motility measured at the widest diameter at 5, 6, 7, 8 and 24 hours. The Sven Gard swarm plates showed more of a swimming phenotype with only the inoculation site showing growth above the agar rather than within for 16/18 of the plates. The other two showed no motility.
