## Supplementary figures and images for "*Salmonella* Genomic Markers for Risk to Food Safety"

### Fig S1

#### Slide 1
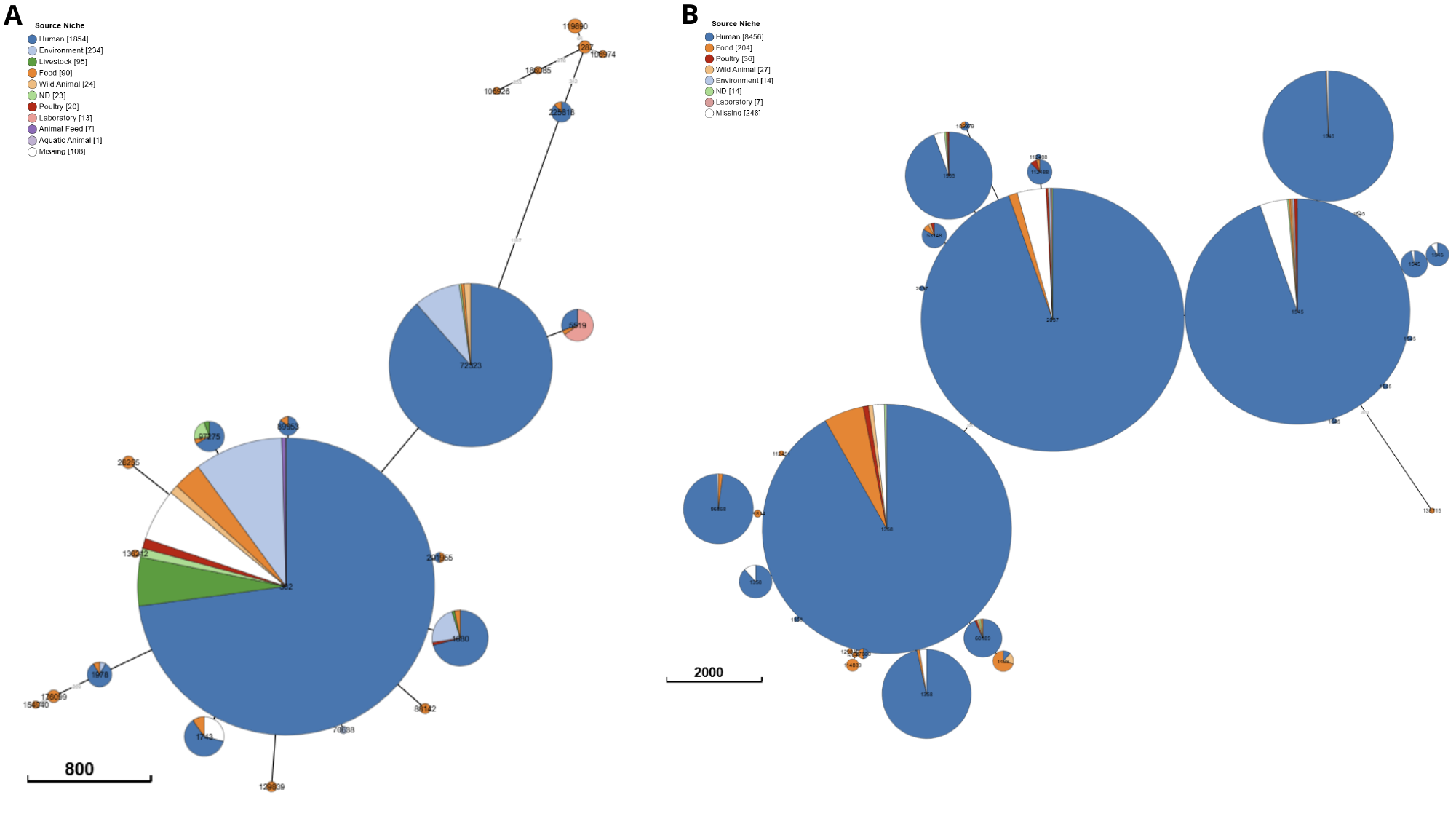

B
A

### Fig S2

#### Slide 1
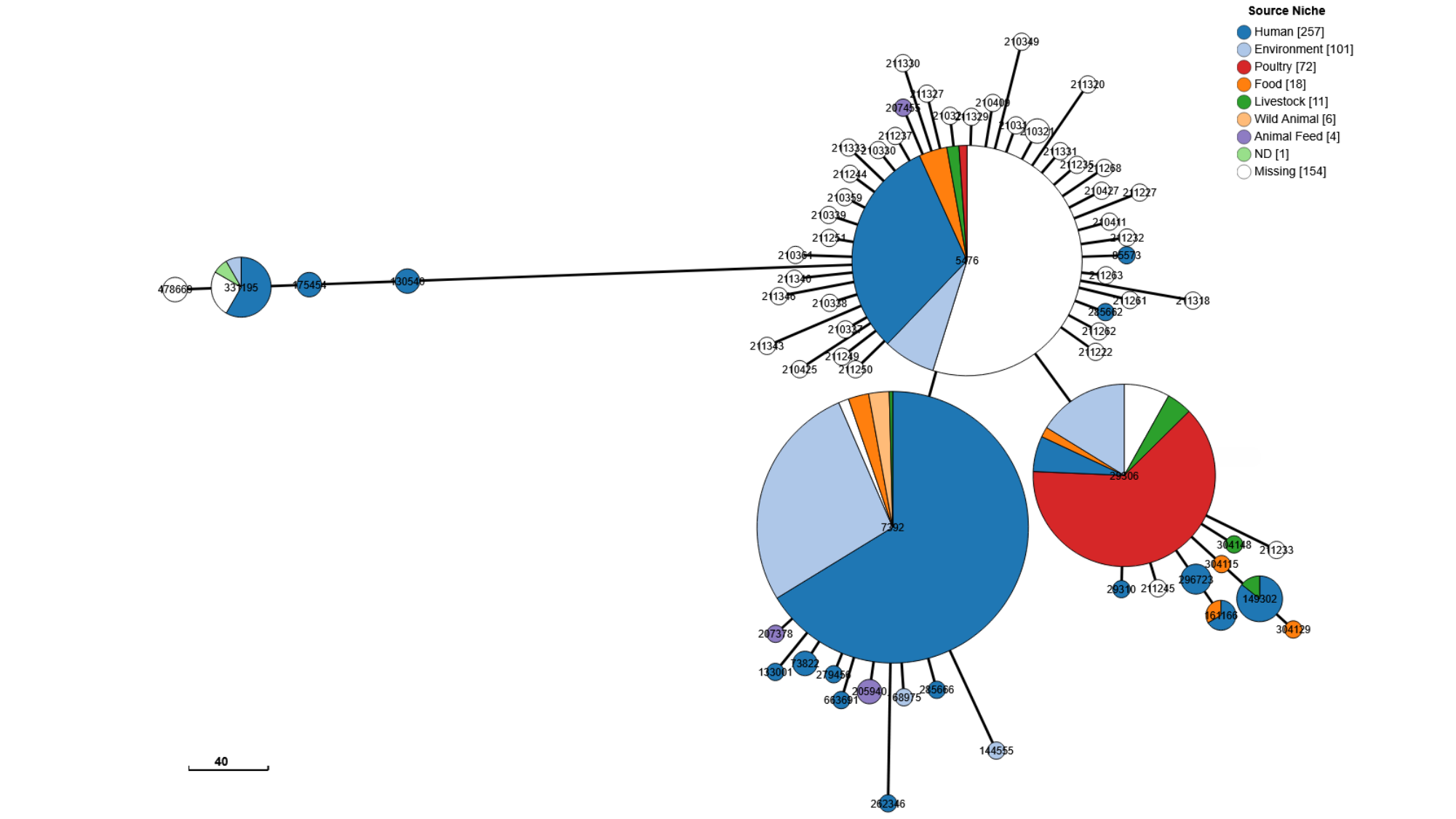

### Fig S3

#### Slide 1
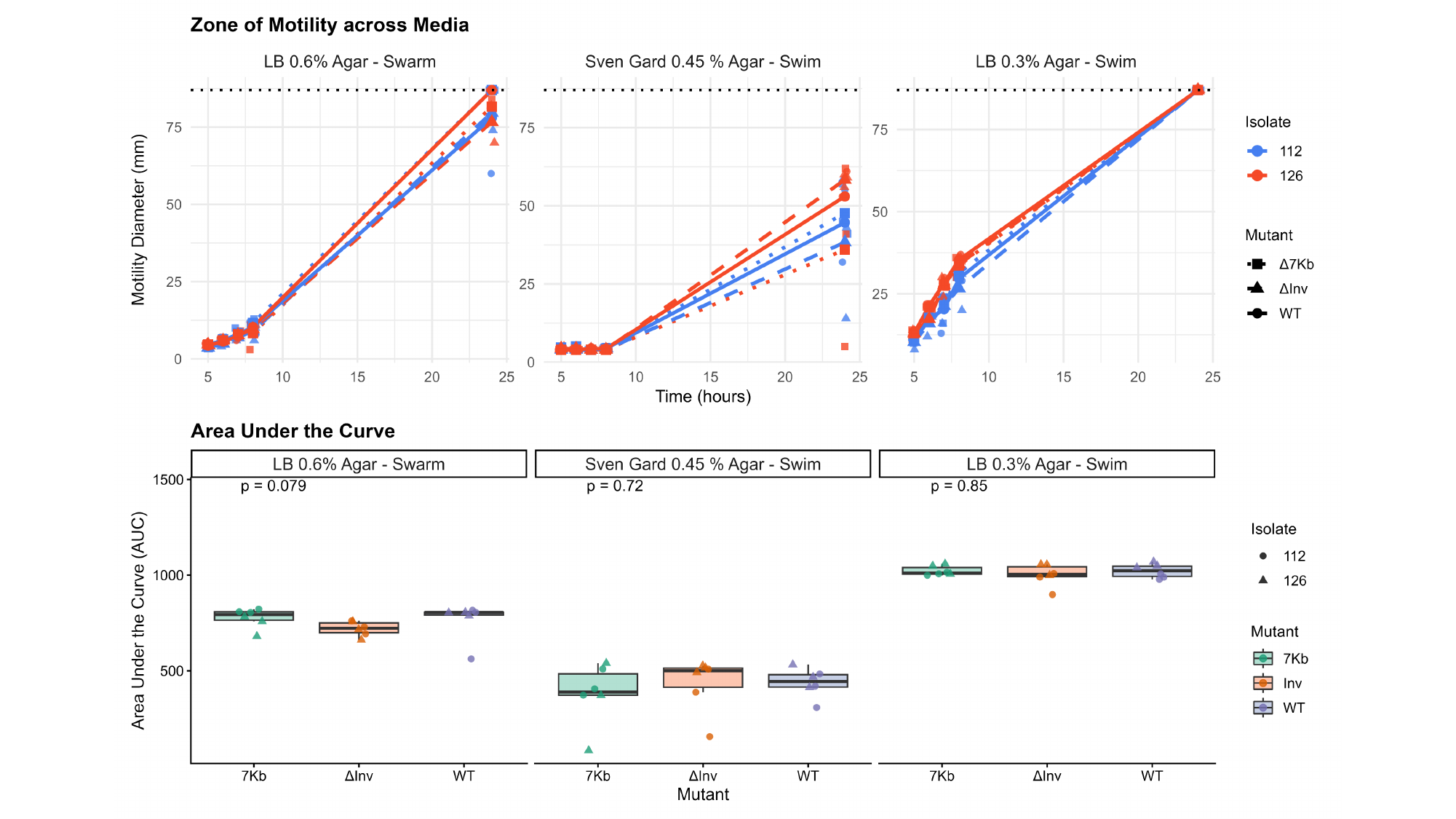
